# PRESTO, a new tool for integrating large-scale -omics data and discovering disease-specific signatures

**DOI:** 10.1101/302604

**Authors:** Sara McArdle, Konrad Buscher, Erik Ehinger, Akula Bala Pramod, Nicole Riley, Klaus Ley

## Abstract

**Background:** Cohesive visualization and interpretation of hyperdimensional, large-scale -omics data is an ongoing challenge, particularly for biologists and clinicians involved in current highly complex sequencing studies. Multivariate studies are often better suited towards non-linear network analysis than differential expression testing. Here, we present PRESTO, a ‘PREdictive Stochastic neighbor embedding Tool for Omics’, which allows unsupervised dimensionality reduction of multivariate data matrices with thousands of subjects or conditions. PRESTO is intuitively integrated into an interactive user interface that helps to visualize the multidimensional patterns in genome-wide transcriptomic data from basic science and clinical studies.

**Results:** PRESTO was tested with multiple input omics’ platforms, including microarray and proteomics from both mouse and human clinical datasets. PRESTO can analyze up to tens of thousands of genes and shows no increase in processing time with a large number of samples or patients. In complex datasets, such as those with multiple time points, several patient groups, or diverse mouse strains, PRESTO outperformed conventional methods. Core co-expressed gene networks were intuitively grouped in clusters, or gates, after dimensionality reduction and remained consistent across users. Networks were identified and assigned to physiological and pathological functions that cannot be gleaned from conventional bioinformatics analyses. PRESTO detected gene networks from the natural variations among mouse macrophages and human blood leukocytes. We applied PRESTO to clinical transcriptomic and proteomic data from large patient cohorts and detected disease-defining signatures in antibody-mediated kidney transplant rejection, renal cell carcinoma, and relapsing acute myeloid leukemia (AML). In AML, PRESTO confirmed a previously described gene signature and found a new signature of 10 genes that is highly predictive of patient outcome.

**Conclusions:** PRESTO offers an important integration of powerful bioinformatics tools with an interactive user interface that increases data analysis accessibility beyond bioinformaticians and ‘coders’. Here, we show that PRESTO out performs conventional methods, such as DE analysis, in multi-dimensional datasets and can identify biologically relevant co-expression gene networks. In paired samples or time points, co-expression networks could be compared for insight into longitudinal regulatory mechanisms. Additionally, PRESTO identified disease-specific signatures in clinical datasets with highly significant diagnostic and prognostic potential.

## Introduction

Large-scale omics data hold great promise to advance precision medicine^1^. Genomic, transcriptomic, proteomic, and metabolomic technologies now provide an unprecedented high-resolution view on all aspects of cell and tissue homeostasis. An integrated, data-driven approach to omics-based health monitoring will broaden our understanding of disease susceptibility and improve diagnostics, treatment, and prevention^2, 3^. Omics technologies measure thousands of parameters per sample, and studies often comprise cohorts with hundreds to thousands of heterogeneous patients.

Excellent tools exist to compare gene^4^ or protein expression^5^ data between two conditions with multiple replicates. However, differential expression (DE) analysis of large data sets with many conditions or confounders and few replicates per group suffers from biological and technical noise as well as low statistical power. Network analysis approaches find groups of markers that are detected in similar patterns across all samples ^6, 7^, independent of experimental groups. This takes advantage of natural biological variation among samples to uncover networks of genes or proteins that are co-expressed under a variety of conditions^8^.

Numerous methods for detecting patterns in –omics data have been proposed and reviewed^9^. Many common approaches for this type of analysis (including principal component analysis^10^, hierarchical clustering^11^, and correlation networks^12^, among others) use various linear metrics to calculate the similarity between genes or proteins. However, these often fail to capture the underlying biological structure^13^, ^14^ because most correlations are non-linear^15^, ^16^. Recently, we described heterogeneity in macrophage polarization by correlating population variability with the expression of known relevant genes^17^. This approach still requires user input (the relevant genes). Weighted gene co-expression network analysis (WGCNA)^18^ uses Pearson correlations weighted by global connectivity to identify robust biological networks. However, WGCNA cannot analyze and visualize the changes in gene networks between conditions with matched samples.

Among non-linear dimensionality reduction techniques, t-stochastic neighbor embedding (t-SNE) has been shown to perform particularly well^19^. It is an adaptation of SNE^20^ that embeds data points in a low number of dimensions in a way that simultaneously preserves both local and global structure from the original high dimensional data. t-SNE has been used for elucidation of cell heterogeneity from single-cell RNA-seq^21,22^, gene networks in Affymetrix arrays^23^, examining similarities between tissues^24, 25^, and unsupervised classification of cells using flow cytometry^26^ and mass cytometry (CyTOF)^27^.

Here, we introduce PRESTO (‘PREdictive Stochastic neighbor embedding Tool for Omics’), a visualization and inference tool for transcriptomic and proteomic data sets. It uses pre-processing filters, t-SNE-based dimensionality reduction, and intuitive visualizations (including movies) for analysis of co-expressed networks of genes. Unlike previous approaches^17^, PRESTO is hypothesis-free and processes all data while blind to sample designation. We demonstrate the usefulness of the tool for exploring co-expressed biological pathways in published pre-clinical data as well as valuable diagnostic signatures from clinical data. A major innovation in PRESTO is the ability to analyze paired data sets to visualize changes in co-expression networks at different timepoints or treatment conditions. PRESTO is provided as a Matlab-based stand-alone tool with an interactive user interface for analyzing -omics data from a variety of experimental designs.

## Results

### PRESTO visualizes co-expressed genes

PRESTO analyzes -omics data from experiments with many samples, conditions, or time points (Fig. 1). The PRESTO algorithm pre-processing filters first select for genes with high variance across all individuals or samples (Supplemental Figure 1A) and sets a minimum expression threshold. The expression and coefficient of variation (CoV) thresholds are user-defined, identifying 1,500-4,000 highly regulated genes. Then, PRESTO utilizes an adapted t-SNE algorithm to map the genes to 2-dimensional space, where dots localized near each other represent genes with similar expression profiles in multi-dimensional space. These scatter plots can be displayed in a variety of ways to visualize expression levels, sample diversity, and changes in co-expression patterns. Density maps are useful for identifying co-regulated networks of genes that are robust and reproducible between investigators. Automated data clustering is an ongoing research area, but no universal “superior” algorithm has emerged yet^28^. The PRESTO user interface includes a module for DBScan^29^ or tSNE coordinate exporting for those users who prefer automated clustering. Gated genes are exported for downstream analysis, functional annotation, diagnostic signatures, and survival predictions. Networks are followed through multiple conditions or time points to visualize changes in co-expression networks. The algorithm is blinded to clinical information, enabling an unsupervised, hypothesis-free analysis of disease phenotypes in thousands of patients or conditions.

**Figure 1:**
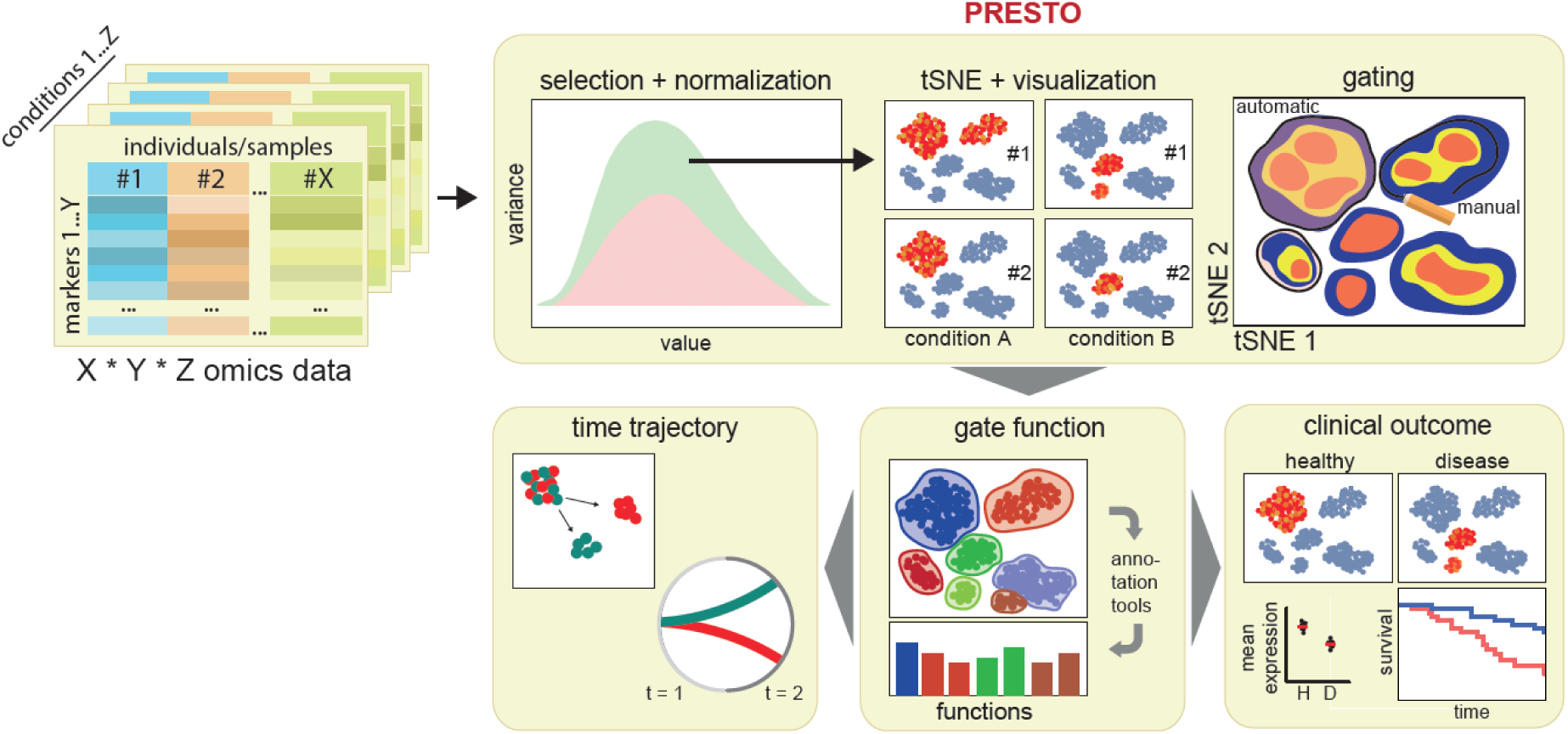
Overview and workflow for PRESTO as a visualization and inference tool for omics data. The input is raw expression data, which is then filtered for markers that are expressed and show significant variance across conditions or subjects. Normalized gene expression is subjected to t-SNE-based dimensionality reduction to find co-expressed networks. Groups of genes are gated and annotated for functions. Time trajectories can be analyzed, for example before and after treatment. Genes in each gate are analyzed for correlation with clinical outcomes like tumor relapseor transplant rejection.

Performance analysis shows that the computing time changes negligibly with the number of columns (i.e. samples) and exponentially with the number of markers (i.e. genes) (Supplemental Fig. 1b,c). We successfully ran an artificial dataset with 100,000 genes (rows), on a high-end commercial desktop PC (data not shown). Moreover, the original t-SNE code^19^ was modified with an automatic exit condition after convergence is achieved. These characteristics make PRESTO uniquely useful for the analysis of large clinical -omics datasets. The package comes as a stand-alone Matlab application (no Matlab license required) with an interactive user interface (Supplemental Fig. 2c).

### PRESTO identifies biologically meaningful gene networks

We first demonstrate the utility of PRESTO using transcriptomes from human peripheral blood mononuclear cells (PBMCs) sequenced via microarray (Supplemental Table 1; 33 samples X 20,898 genes; GSE74816)^30^. The PRESTO pre-processing filters (Supplemental Table 1) retained 1,298 genes. tSNE-based dimensionality reduction organized the genes into 9 dense networks (Fig. 2a,b). Each of these gates contains a set of genes whose expression across the 33 samples tends to rise and fall together, relative to their individual average expression. PRESTO ranked relative expression across all 33 patients for each gene independently by coloring the dots from highest (red) to lowest (blue) (Fig. 2c, Supplemental Fig. 2). The genes that were not gated with others are those that vary between subjects but not in a substantially similar way to other genes that were mapped. The genes in each gate (Fig. 2b) were significantly enriched for various functions (Fig. 2d), for example B cell activity (gate 1) and Fc and complement signaling (gate 2). All functional enrichments are shown in Supplemental Table 2.

**Figure 2.**
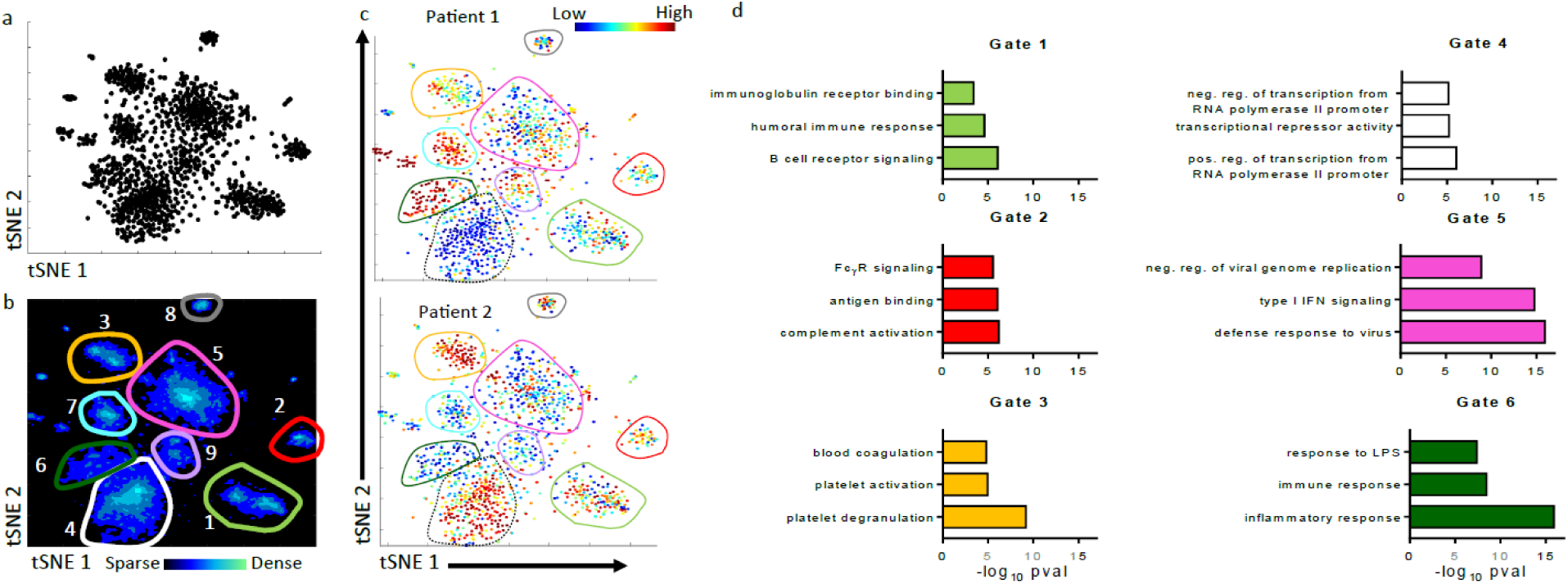
PRESTO identifies co-expressed gene networks in healthy human PBMC transcriptomes. Gene expression in PBMCs from 33 patients were analyzed with PRESTO (GSE74816). Pre-processing filters (CoV threshold = 2) retained 1,298 genes. A) Dimensionality reduction organized the genes into 2 dimensions. B) A density map of the 2D output identifies 9 major gates. C) Relative expression plot from 2 subjects. For each gene (each dot), the 33 samples were ranked from the lowest (blue) to highest expressers (red). Supplemental Fig. 2 shows the other subjects. D) Functional annotations for 6 of the gates were found using DAVID. p-values for gene enrichment by modified Fisher’s exact test.

DBScan clustering produces results very similar to density map gating (Supplemental Fig. 3a,b). Attempting to discover groupings in the 1,298 PBMC genes by PCA produced no obvious pattern and hierarchical clustering detected groups that showed little similarity with PRESTO gates (Supplemental Fig. 3c,d). K-means clustering of the tSNE map failed to detect the visible boundaries between the gates (Supplemental Fig. 3e). K-means clustering of the 33-dimensional raw data also showed no pattern in tSNE space (Supplemental Fig. 3f). WGCNA produced colored bars that identified groups of genes similar to PRESTO gating (Supplemental Fig. 3g,h).

The patterns derived from PRESTO are insensitive to changes in the input data or user-controlled settings. Changing the randomly generated initial seed coordinates, the proportion of genes included, the number of subjects included, or the “perplexity” settings (which determines the effective number of nearest neighbors)^19^ made little difference to the final results (Supplemental Fig. 4a-e). Raising the CoV threshold sharpens the boundaries between the gates, but notably, the original gates remain separated under all conditions (Supplemental Fig. 4f).

### PRESTO identifies gene networks in the LPS response of mouse peritoneal macrophages

To show PRESTO’s ability to detect co-expressed networks in populations with high levels of natural variability, we used microarray data of peritoneal macrophages harvested from 75 inbred mouse strains in the Hybrid Mouse Diversity Panel (HMDP) treated with lipopolysaccharide (LPS) (Supplemental Table 1; 75 strains x 13,699 genes; GSE38705)^31^. PRESTO pre-processing selected 2,423 genes that were gated into blue, green and red gates based on density maps (Fig. 3a). Functional annotation using DAVID^32^ showed the green genes were enriched for inflammation, the blue genes for chromatin organization, and the red genes for cell cycle (Fig. 3b) (p<.0001 to <.05). PRESTO’s relative ranking function (Fig. 3c, Supplemental Movie 1, Supplemental Fig. 5, Supplemental Table 3) made diverse macrophage polarization among mouse strains immediately and intuitively obvious.

**Figure 3.**
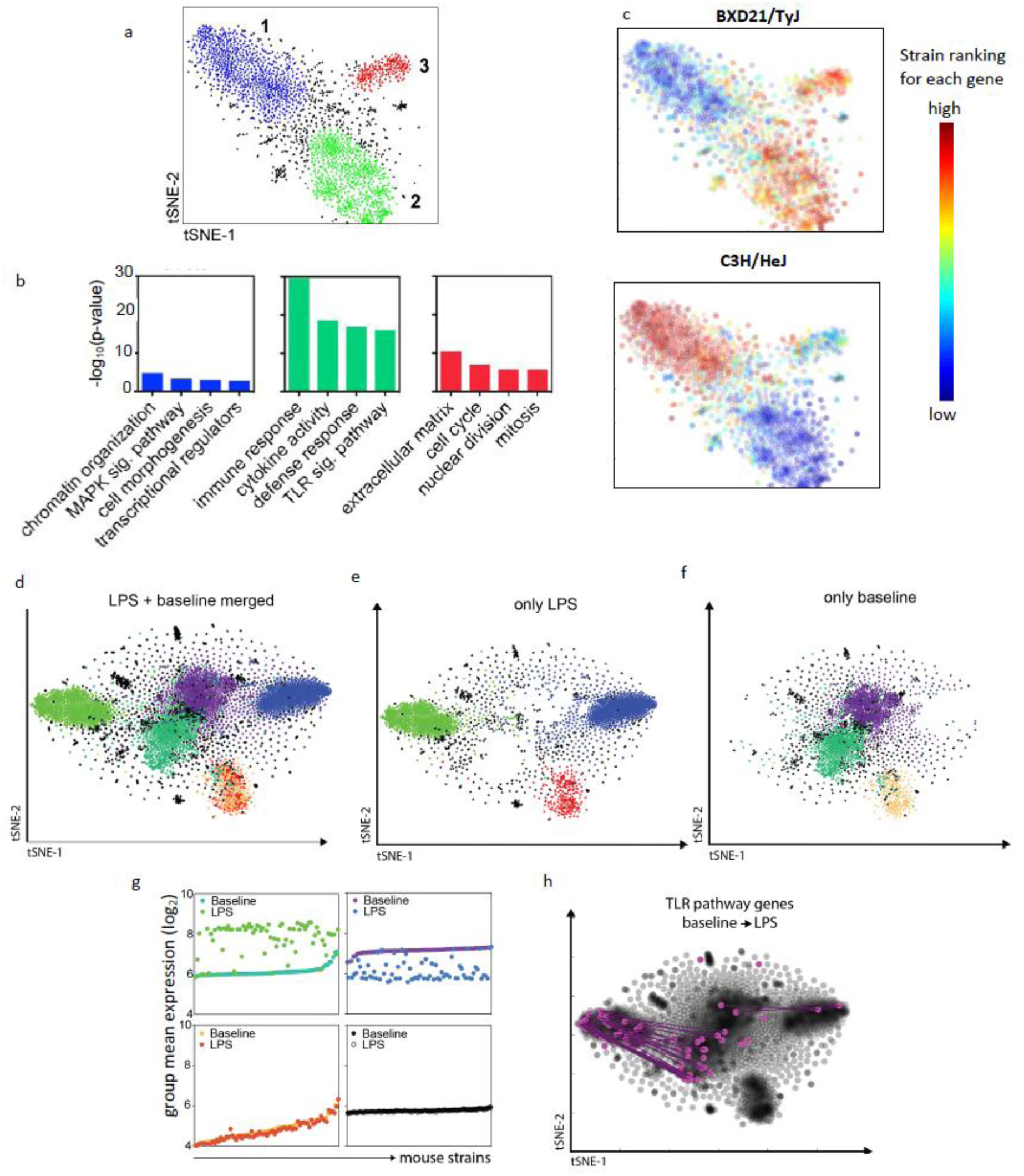
PRESTO identifies LPS-induced movement of three distinct gene networks in peritoneal macrophages of 75 mouse strains. Transcriptomes (GSE38705) from peritoneal macrophages harvested from 75 inbred mouse strains treated with LPS were filtered (to 2,423 high-variance genes) by PRESTO and automatically organized (a) into 3 major gates (CoV threshold = 1.5) b) Selected DAVID annotations of the 3 PRESTO gates and their enrichment p-values (modified Fisher’s exact test). c) Each mouse strain was ranked from highest (red) to lowest (blue) expression of each gene and the ordinal ranking is plotted for each gene (dot). Strains BXD21/TyJ and C3H/HeJ are shown here. See Supplemental Figure 5 and Supplemental Movie 1 for all strains. d-h) The transcriptomes of LPS-treated and untreated peritoneal macrophages from 75 inbred mouse strains were analyzed concurrently by PRESTO in an unsupervised and blinded manner. Resulting scatter plots of d) the combined map and e-f) separated baseline and LPS transcriptomes. Each gene has a color-coded designation based on the gates in a. The purple and cold green gates at baseline move to the blue and warm green gates after LPS (Supplemental Movie 2). The genes in the orange gate (red in LPS-treated) do not move (LPS-unresponsive genes). g) The mean expression of all genes in each gate was calculated for each strain, comparing the LPS-treated and untreated expression of each gate. Mouse strains were ordered in ascending order of average baseline expression in each group. Expression after LPS is shown as a dot above or below. h) Known TLR pathway genes are highlighted to show the movement of these genes from the untreated to the LPS-stimulated state. Most TLR pathway genes are in the cold to warm green gates. Vectors connect each gene at baseline and after LPS.

We repeated the robustness tests on the LPS-treated macrophage dataset and found the resulting scatter plot to be insensitive to changes in user-defined settings, random initial seeds, or small changes in the input data, while conventional methods underperformed (Supplemental Figs. 6 and 7).

To explore LPS-induced changes in co-expressed gene networks across the 75 mouse strains, we combined the LPS-treated transcriptome with that from untreated peritoneal macrophages (Fig. 3d, Supplemental Table 1; 2 conditions x 75 strains x 13,699 genes; GSE38705)^31^. Genes from both conditions were paired by mouse strain, and the same 2,423 genes from both data sets were analyzed. Gates were defined on the LPS-treated data set (Fig. 3a). The off-color scheme of purple, cold green, and orange in baseline corresponds to blue, red, and warm green after LPS (Fig. 3d-g). Between baseline and LPS, the blue genes move to the right in tSNE-1, the green genes move to the left, and the red genes do not move (Supplemental Movie 2). Remarkably, all gated genes remained together, suggesting that these gene lists form networks both before and after LPS stimulation. On average, genes in the green gate are upregulated and those in the blue gates are downregulated by LPS (Fig. 3g). The large majority of genes known to be part of the Toll-like receptor 4 (TLR4) pathway (the major receptor for LPS in macrophages^33^) that pass the pre-processing filters are in the green, inflammation-related gate (Fig. 3h). In contrast, the genes in the red gate stayed together before and after LPS stimulation and did not move. This suggests a consistent pattern of expression across the 75 strains that was not influenced by LPS. The unlabeled (black) genes were not gated, suggesting that there is no regular high-dimensional pattern linking them. This can be further seen in the gate expression means, where the red genes vary considerably across strains but are unchanged by LPS, while the black-labeled genes are flat across strains (Fig. 3g).

### PRESTO finds gene networks across the human macrophage activation spectrum

To demonstrate the ability of PRESTO to visualize patterns in data with many conditions (instead of many subjects or strains), we analyzed transcriptomes of cultured human monocyte-derived macrophages treated with different stimuli (33 conditions x 1 subject x 15,798 genes; GSE68854)^34^. These treatments (Supplemental Table 1) are known to induce different macrophage phenotypes; i.e. LPS and interferon-gamma (IFNγ) induce classically activated macrophages, while Interleukin-4 (IL-4) induces alternatively activated macrophages^34, 35^. PRESTO-based density maps identified 8 gates (Fig. 4a), which contain genes with distinct biological functions (Fig. 4b). Interestingly, gate 1 (138 genes) was clearly delineated in LPS, IL-4, immune complexes (IC), IL-10, IFNβ and IFNγ, but not in GM-CSF or dexamethasone-treated macrophages (Fig. 4c). However, gate 1 did not contain many annotated genes. This suggests that PRESTO can identify new gene networks with unknown functions and pathways. As exemplified by 8 classical activators (of 33 in the data set), each stimulus induces a characteristic pattern of up- or downregulated gene networks (Fig. 4c). For example, the genes in gate 3 are only highly expressed upon LPS, GM-CSF, or IFNγ treatment. To more formally analyze this, we used hierarchical clustering of the mean expression of all genes in each gate (Fig. 4d). The known classical macrophage inducers GM-CSF, LPS, and IFNγ cluster together, as do the known alternative stimuli dexamethasone, IL-4, IL-10, and IC.

**Figure 4:**
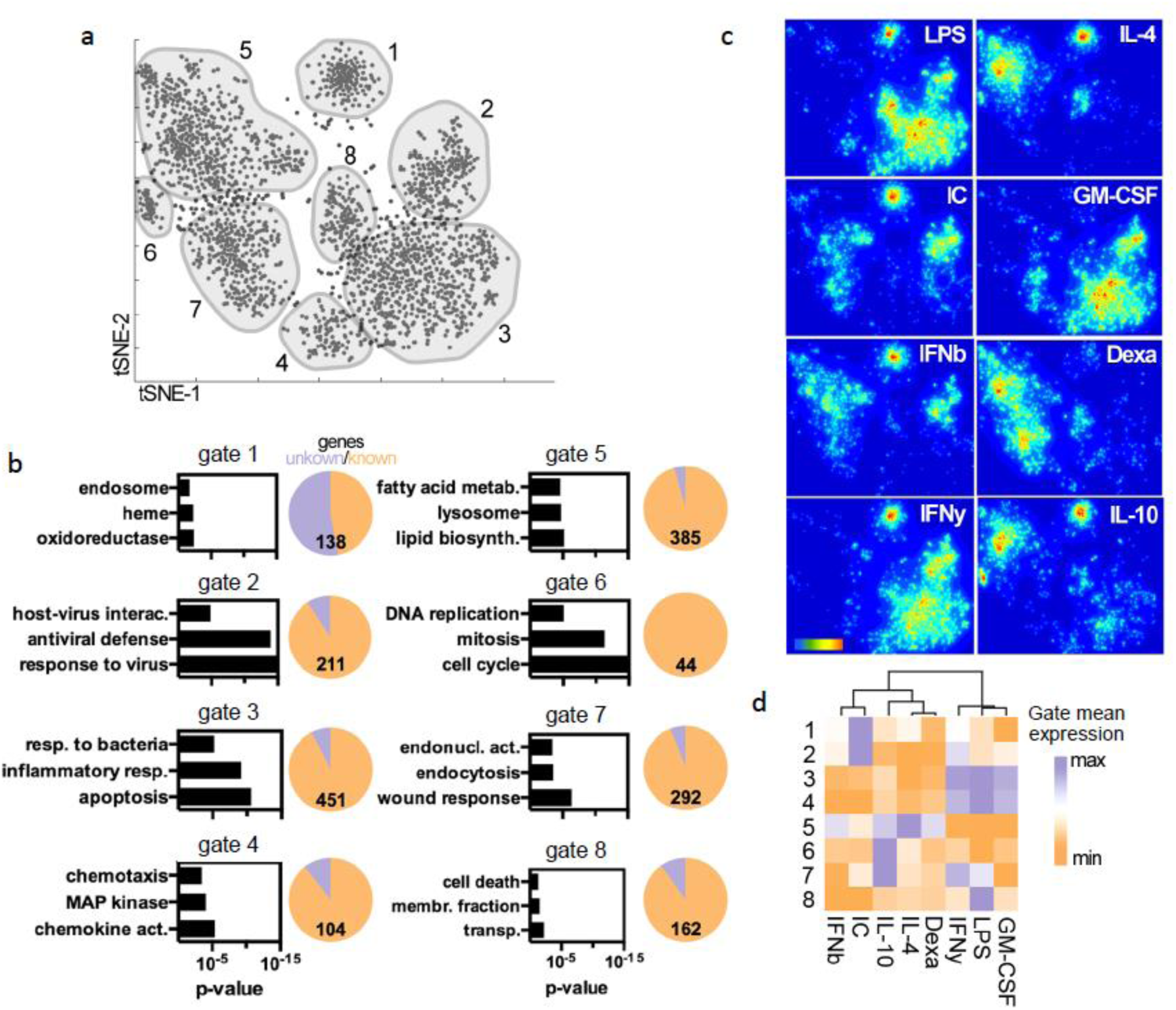
Stimulus-induced gene networks in human macrophage activation. Human monocyte-derived macrophages from one donor were treated with 33 different stimuli *in vitro*, and analyzed with RNA microarray (GSE68854). a) PRESTO selected 2,083 genes and organized them into 8 distinct gates. b) Selected DAVID annotations for the depicted gates and their enrichment p-value. The pie charts show the total number of genes and the fraction with no annotation (=unknown) in each gate. c) For each of 8 canonical stimuli, the genes that are expressed highly in that sample compared to other stimuli (similar to red dots in Figs. 2c and 3c) are isolated and shown as density plots to reveal selective activation of gene networks. d) Gene gate means for the 8 stimuli were hierarchically clustered to find stimuli that induce similar expression profiles.

### PRESTO determines co-expression network changes in human vaccine response

The PBMC transcriptome (Fig. 2) was collected as part of a larger study into the response to influenza vaccine, in which PBMCs were isolated at days 0, 3, and 7 after vaccination (Supplemental Table 1; 3 time points x 33 patients x 20,898 genes). Pre-processing filters selected the same 1,875 genes from each of the 3 matched time points. These were mapped onto the same tSNE axes, blinded to both gene and sample identity (Fig. 5a). 19 gates were drawn based on the density map. Dots representing the genes from day 0, 3 and 7 were then separated to show the gene networks present on each day (Fig. 5b, Supplemental Movie 3). Circos plots^36^ revealed that some genes stayed in the same gate, suggesting consistency across time, while others moved to a different gate or even split to populate 2 or more gates. The blue gene network (nucleosome and histone) in gate 14 at day 0 moves to gate 15 on day 3 and to gate 2 on day 7 (Fig. 5d). This suggests that this network of genes is regulated by a similar mechanism across subjects, but regulation changes over time. The purple gene network in gate 1 on day 0 stays in gate 1 on day 3 and then splits to populate gate 4 and ungated (gate 0) on day 7 (Fig. 5e). This suggests that these genes are co-expressed at day 0 together with day 3, but not day 7. Gate 1 is enriched for genes involved in B cell activation, which is known to take place approximately a week after a stimulus. The yellow gene network (inflammatory response and chemotaxis) is not well developed on day 0, appears in gate 17 on day 3 and then splits into gate 6 and 10 on day 7 (Fig. 5f). This means that some of the inflammation and chemotaxis genes are initially co-regulated with each other and then co-regulated with other gene networks. This loss of co-expression after a stimulus could have important implications on the human variability of response to vaccines. This kind of analysis allows visualization (Supplemental Movie 3) of how gene networks are rearranged over time, providing insight into regulatory mechanisms.

**Figure 5.**
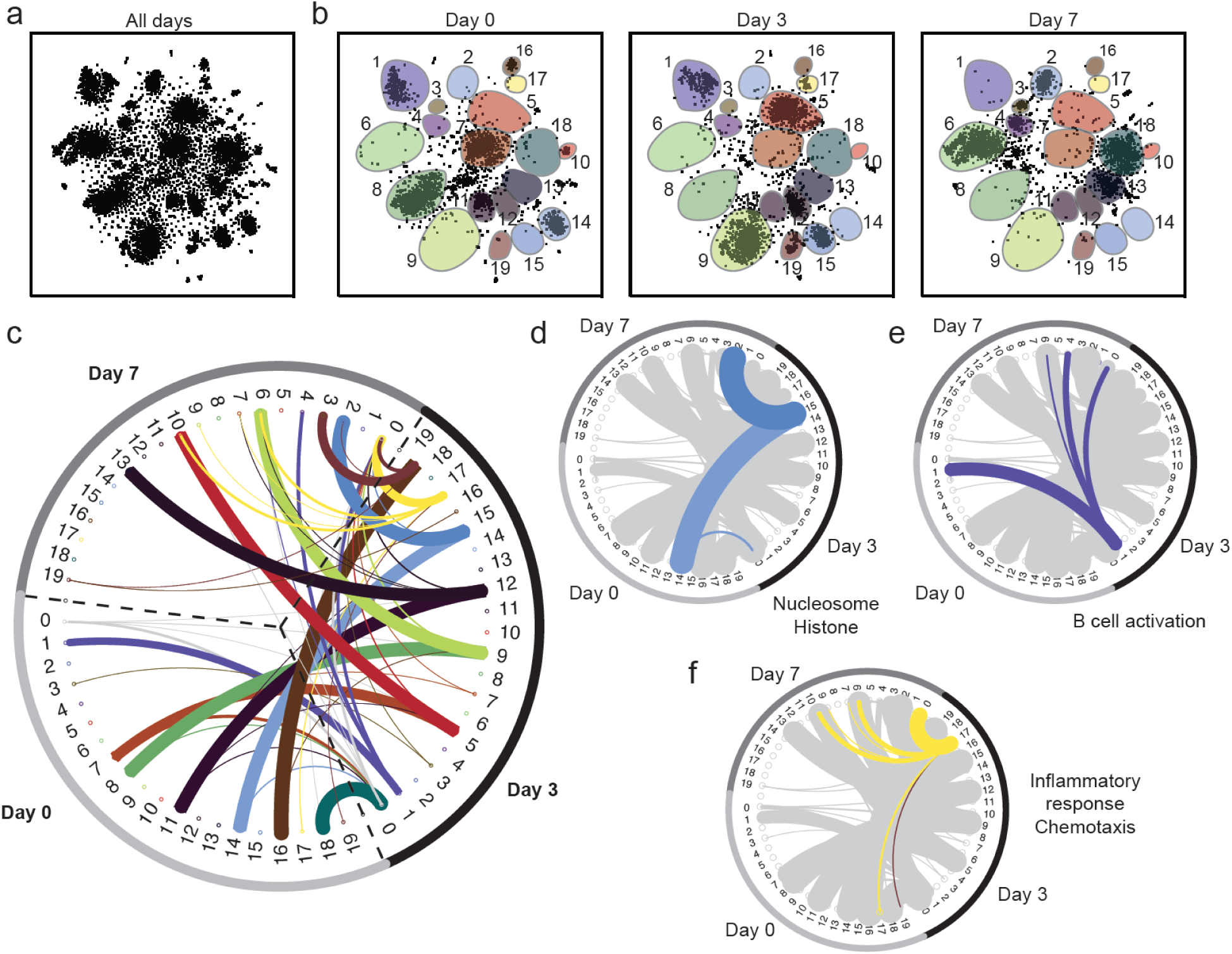
Gene network coherence over time after influenza vaccination. PBMC transcriptomes from 33 human subjects at days 0, 3, and 7 after flu vaccination (GSE74816) were analyzed with PRESTO, keeping each gene 3 times (once for each time point). a) 1,875 genes passed the pre-processing filters for each time point, leading to 5,625 data points mapped onto the same axes. b) After mapping, the genes from each day are separated to display changes in the co-expression patterns with time. Gates found by density maps (not shown) superimposed. c) Circos plots show how the genes transition between gates at 0, 3 and 7 days. The width of each line shows how many genes make any given movement, normalized to the total number of genes found in a gate (Supplemental Movie 3). Ungated genes are assigned to group 0. For clarity, any transition of fewer than 2% of the total genes found in a gate was removed. e-f) Representative transition patterns are highlighted for clarity. Enriched functional pathways for each are shown (p<.01, modified Fisher’s exact test.

### Prognostic gene signatures in human kidney transplant rejection

Tissue biopsies followed by histological assessment are the gold standard for diagnosis of solid organ transplant rejection. Biopsy sequencing is a promising method of increasing sensitivity of diagnosis^37^. We applied PRESTO to renal biopsy transcriptomes of 48 patients with healthy or rejecting (antibody-mediated rejection, AMR) kidney transplants (Supplemental Table 1; 2 conditions x 48 patients x 20,647 genes; GSE50084)^38^. Density plots identified four major gates (Fig. 6a). Mean expressions of the genes in gate 1 across all patients revealed a rejection-associated signature with strong upregulation of expression (Fig. 6b), enriched for inflammation, immunity, and allograft rejection (Fig. 6c). Gates 2 - 4 were enriched for basal transport and cytoskeletal processes (data not shown). The previously published DE analysis showed a total of 2,354 genes significantly upregulated in rejected transplant biopsies^39^. PRESTO identified an inflammation-enriched gene signature of 388 genes (gate 1).

**Figure 6:**
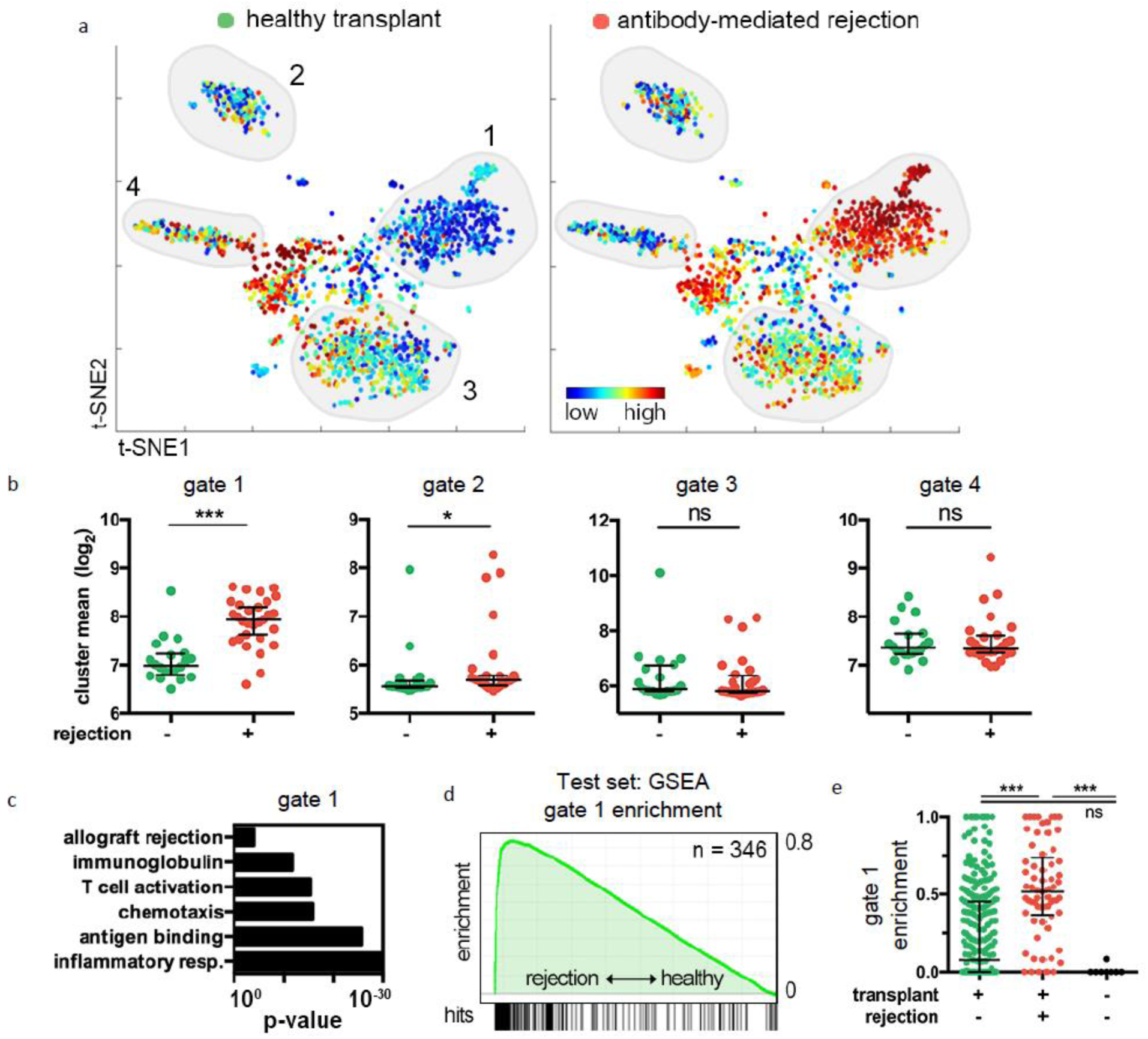
Gene signatures from kidney biopsies indicate kidney transplant rejection. 20,873 genes from kidney biopsies from patients with healthy transplants (n=20) or antibody-mediated rejecting (AMR, n=28, GSE50084) kidney transplants were analyzed. PRESTO was blinded to all clinical data. a) Relative ranking maps of a representative healthy (left) and representative rejecting (right) individual. PRESTO selected 1,549 genes (CoV>2.1) and organized them into 4 gates identified by density plots (not shown). For each dot, blue (low) and red (high) colors indicate the relative expression of that gene in an ordinal scale across all patients. b) Gate mean expression for all individual patients. The classification of non-rejection or rejection is based on histological assessment. Data shown as gate median +/- IQR. Two-tailed Mann-Whitney test. ^∗∗∗^, p<.001; ^∗^, p<.05 c) Selected DAVID annotations for gate 1 and their enrichment p-value. d,e) Independent test data (GSE36049) set of healthy (n = 281) and rejecting (n = 65, AMR) kidney transplant biopsies. e) Gene-set enrichment analysis with significant enrichment of gate 1 genes only in rejection but not healthy biopsies. FDR < 25%. f) Deep RNA deconvolution of raw biopsy transcriptomes based on the group 1 gene profile successfully identifies rejection patients. Controls included healthy nephrectomies (non-transplanted). Median +/- IQR indicated. Kruskal-Wallis test with Dunn’s multiple comparison test.

To verify the diagnostic robustness of PRESTO signatures, we used the genes from the PRESTO gates to detect allograft rejection in an independent test set^40^ (GSE36049) of 346 non-rejecting or AMR kidney transplant patients (Fig. 6d). Using gene set enrichment analysis (GSEA), genes in gate 1, but not gates 2 – 4, were found to be highly enriched in rejection (Fig. 6e, and data not shown). Similarly, RNA deconvolution successfully detected the gate 1 rejection signature in these biopsies and discriminated between healthy and rejection samples (Fig. 6e). The gate 1 genes were not enriched in native (non-transplanted) kidney controls (samples derived from nephrectomies). These data suggest that PRESTO readily identifies a robust rejection-specific gene signature in large clinical cohorts using bulk biopsy transcriptomes.

As a proof-of-concept that PRESTO is applicable to other types of -omics data, we analyzed published data of label-free mass spectrometry from clear cell renal cell carcinoma (ccRCC) biopsies taken from 84 patients with stage 1-4 tumors or adjacent tumor-free tissue (Supplemental Table 1; 2 conditions x 84 patients x 783 proteins)^41^. Four distinct protein gates were identified (Supplemental Fig. 8a). Gate 3 was highly specific for proteins involved in tumor-related processes (Supplemental Fig 8b), and significantly upregulated in all tumor stages (Supplemental Figure 8c and data not shown). Using this gate as a gene list, many master regulators known to affect renal cell carcinoma malignancy were predicted by Ingenuity Pathway Analysis, including the tumor suppressor gene VHL and the associated factors HIF1A and VEGF (Supplemental Figure 8d)^42^. This demonstrates PRESTO’s versatility and applicability across data formats and -omics platforms.

### Gene signatures to predict leukemia relapse and patient survival

Acute myeloid leukemia (AML) still has poor outcomes, and predictive gene signatures that could guide therapeutic decisions are sorely needed. Here, we tested whether PRESTO can derive clinically useful gene signatures from publicly available AML datasets. PBMCS were sorted into cell fractions that were characterized as containing stem cell activity (LSC+) or not (LSC-)^43^ (Supplemental Table 1; 227 cell fractions x 19,529 genes; GSE76008). Ng, et. al. found that a signature of 17 genes associated with stemness was predictive of AML outcome.^43^

PRESTO is inherently hypothesis-free. Therefore, we tested whether PRESTO would automatically identify a predictive gene signature containing the stemness genes, without prior knowledge of these genes or even without knowing sample stratification. We reasoned that PRESTO might identify other genes that contribute to predicting AML outcome. PRESTO identified 6 gates (Fig. 7a). The ratio of expression between LSC+ and LSC-samples for each gene was calculated post-hoc. Gate 5 (orange) is enriched for genes that are higher in LSC+ samples (Fig. 7b). However, a gene signature of 584 genes is too large to be clinically useful. Therefore, we reapplied PRESTO to the genes in gate 5, which further divided the genes into 6 new gates (Fig. 7c). Gate A contains the genes expressed most highly by LSC+ samples. Gate A genes (48 out of 50) were successfully matched to an independent test data set of PBMC transcriptomes from 160 AML patients (GSE12417). A hazard signature (Sig A, Table 1) was calculated with Cox proportional hazard regression (CPHR). Figure 7e shows that patients with an above-median hazard score have significantly shorter survival (p<0.00001, hazard ratio (HR)=3.7). To refine the predictive signature, we removed the genes that contributed little to the survival prediction (those with p>.1 based on CPHR), resulting in a condensed signature of 13 genes (Signature B). CPHR coefficients recalculated with only this list (Signature B) still significantly predicts outcome (p<.00001, HR=4.3). Signature B contains 3 of the 17 known stemness genes^43^. Therefore, we asked whether the known stemness genes were driving the prediction, or whether the other 10 PRESTO-discovered genes were sufficient to predict survival. Removal of the stemness genes (SOCS2, BEX3, and CPXM1) leads to a unique 10-gene (Signature C), which still predicts survival at the same level of significance. 9 of these genes have been described in AML or other cancers (Table 1)^44-53^. One gene, SHANK3, is completely novel. This shows that PRESTO can identify new clinically useful gene signatures that predict survival in AML patients.

**Figure 7:**
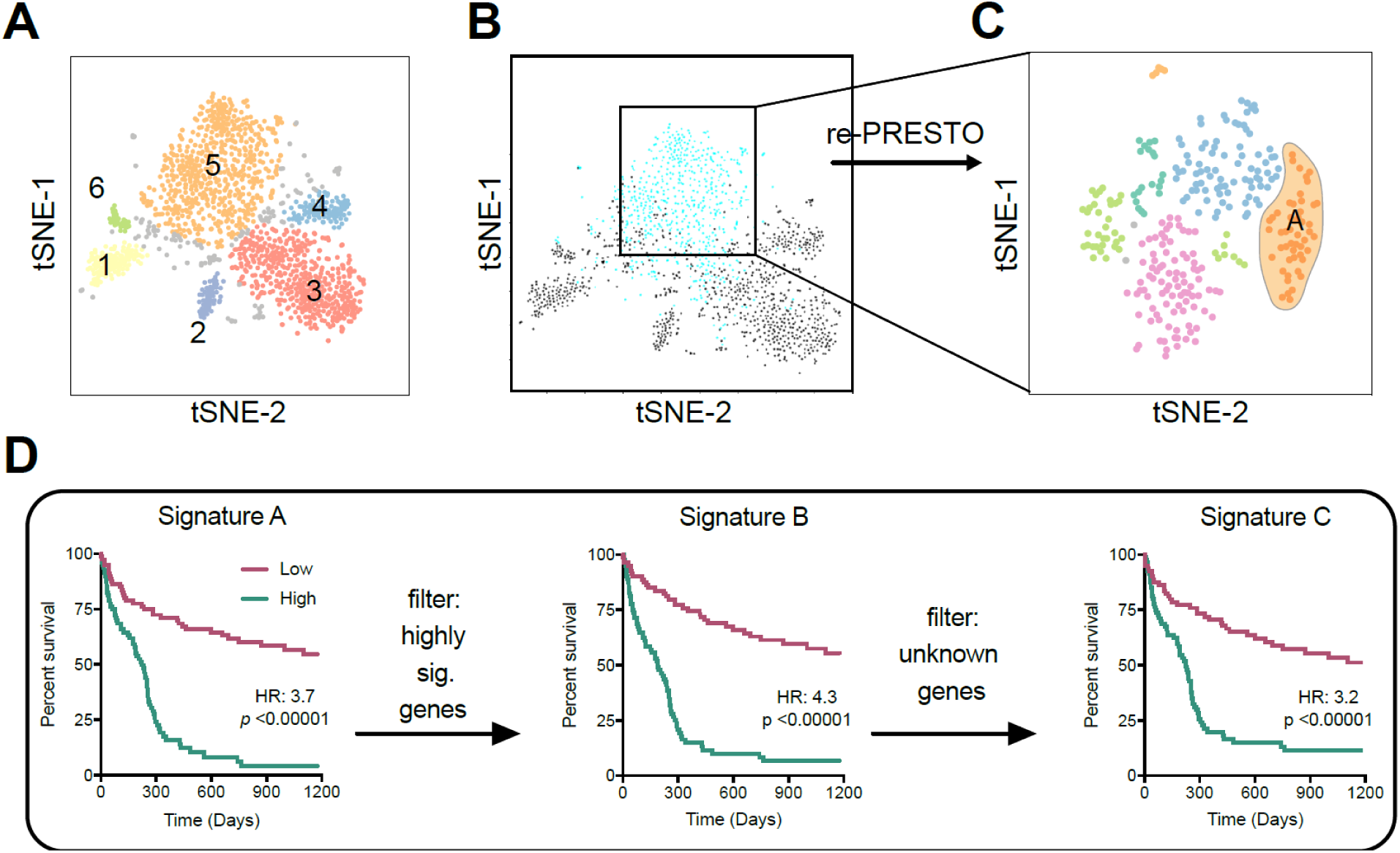
PRESTO identifies gene signatures that predict acute myeloid leukemia (AML) survival. A)Transcriptomes from 227 sorted PBMC fractions (GSE76008) taken from patients with AML were analyzed with PRESTO. Gates based on density maps (not shown). B) The ratio of expression in the LSC+ samples compared to the LSC-samples was calculated, and the genes with a ratio>1 are shown in blue. C) The genes in gate 5 were analyzed more deeply through a second round of PRESTO applied to gate 5 only. The 584 genes were filtered to 290 extremely variable genes, and after dimensionality reduction 7 gates were identified. D) 48 of the Group A genes could be matched to a test data set comprised of PBMC transcriptomes from 163 AML patients (GSE12417). These genes were analyzed with Cox proportional hazard regression (CPHR) to create Signature A. Kaplan-Meier curves show that patients with the above-median Signature A score have significantly worse survival than the other half of patients in the test data set. Removing the genes which minimally contributed to the survival prediction (those with p>.1 based on CPHR) leaves a condensed list of 13 genes. Re-calculating the CPHR coefficients with only this list generates Signature B, which significantly predicts survival in the test data set. Further removing genes that have been previously reported to predict AML survival^43^ leads to 10-gene Signature C with highly significant predictive power.

**Table 1.**
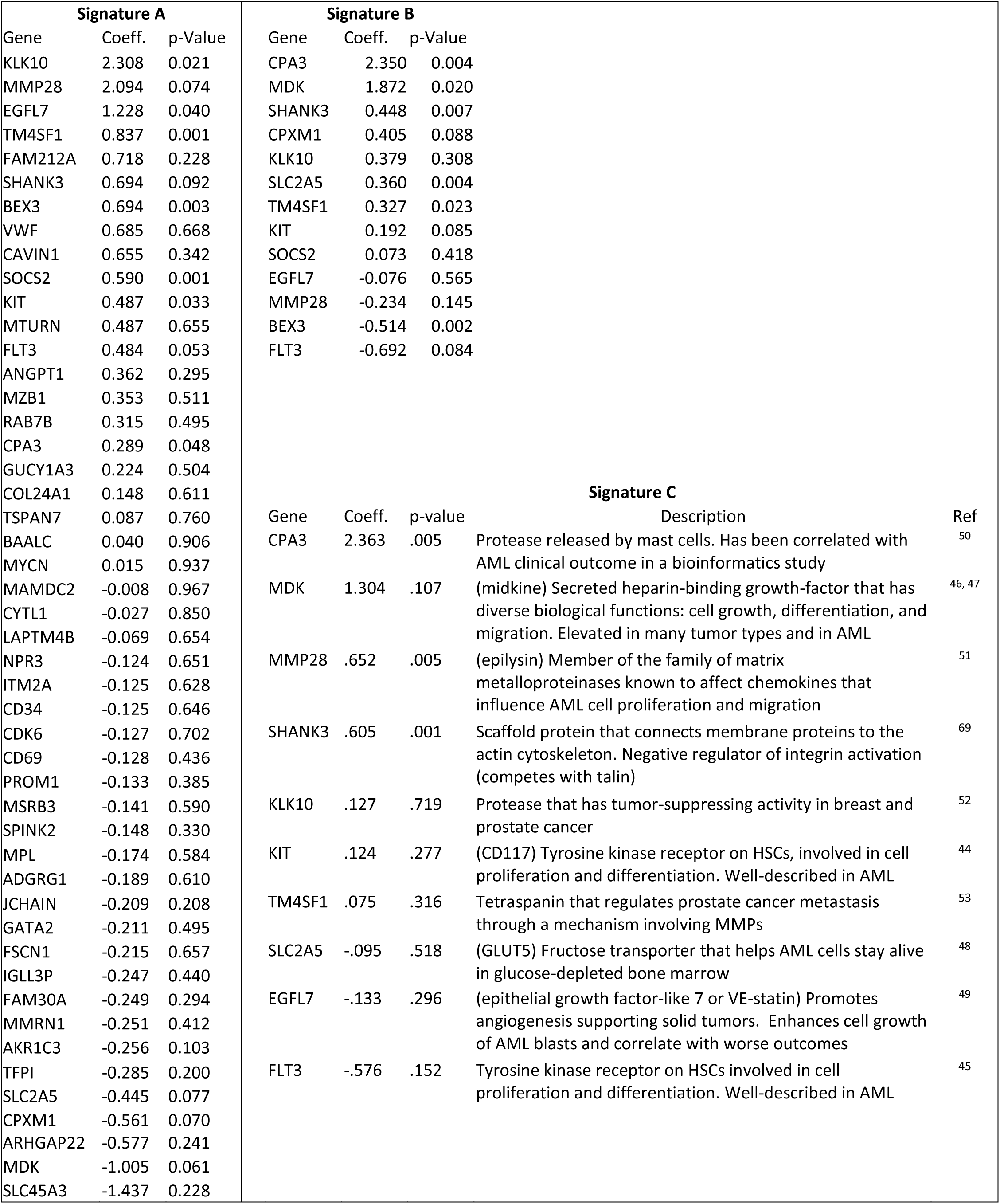
CPHR coefficients and p-values for the genes in Signatures A, B, and C. Known functions are provided for the genes in Signature C.

## Discussion

We introduce PRESTO as a tool for co-expression analysis and visualization of -omics data, with several prominent features: 1) it organizes genes with similar expression profiles more clearly than PCA or heat maps, 2) it performs well on -omics data derived from clinical samples (bulk biopsies), 3) it processes very large datasets (20,000 markers x 100,000 samples) on standard commercial computers, 4) it provides multiple intuitive visualization and output options for downstream analysis, 5) it natively processes paired data sets for simultaneous comparison of multiple conditions, 6) it can handle different -omic data types, and 7) it generates highly reproducible results. In the age of high-throughput technologies, these attributes render PRESTO uniquely useful for many large-scale studies in precision medicine.

In most bioinformatics applications involving dimensionality reduction, the displayed data points are individual cells or samples^21,22, 24-27^. In contrast, PRESTO analyzes the markers themselves, looking for patterns in expression across all of the varied samples in a data set. While t-SNE based methods have been used to study gene co-expression networks previously^23^, the previous approach required a pre-defined list of DE genes. This severely limits the applications of the method to only those where DE analysis is relevant, whereas PRESTO is hypothesis-free and does not require DE analysis.

A main benefit of PRESTO is that it is applicable to a wide variety of experimental designs, including those with multiple groups, many confounders, and low numbers of replicates. Furthermore, unlike WGCNA, PRESTO analyzes paired samples (time series, different culture conditions of the same cell types, etc.). This is possible because the marker names are the same between different experimental groups, allowing them to be tracked. Paired projection onto the same axes simultaneously allows for direct comparison of co-expressed networks in multiple conditions. The change in gene location can be shown as a movie to visualize alterations in co-expressed networks (Supplemental Movies 2,3). While the distance between gates on a tSNE plot is difficult to interpret, the merging or splitting of gene networks is suggestive of biologically meaningful changes to regulatory mechanisms.

The most relevant way to evaluate a transcriptomic analysis algorithm is to interrogate the gene list(s) it produces^13^. DAVID identified enrichment of known biological functions in many of the gene networks that reflect the context of the experimental or clinical conditions, underlining PRESTO’s usefulness. Additionally, PRESTO grouped genes with unknown functions together with well described pathways, suggesting a starting point for investigation into the functions of these genes^7^.

We demonstrate that PRESTO-derived gene signatures from human transcriptomic and proteomic data sets might be clinically useful for diagnosis and survival prediction. PRESTO analysis is unsupervised, bottom-up, and therefore, highly advantageous for hypothesis-free discovery applications. Patient stratification is correlated with the gates post-hoc for unbiased discovery of disease specific networks. Applying a second round of PRESTO to a gate enables condensing hundreds of genes to a smaller, specific signature that is more feasible for clinical routine^54^.

As proof-of-concept, we analyzed a clinical data set of AML in depth and found a new signature (10 genes) that significantly predicts survival time. These genes had not been reported by the previous study using the same training data^43^, establishing that PRESTO is complementary to other bioinformatics techniques. Separate studies described nine of the 10 genes in this signature as being clinically correlated to cancer pathophysiology or prognosis^44-53^. PRESTO was able to discover known AML-related genes even though it was blinded to the sample designation, demonstrating biological relevance of the technique. Furthermore, it identified a novel gene, SHANK3, as relevant to AML. SHANK3 is scaffold protein that connects membrane proteins to the actin cytoskeleton. As of the time of this writing, we could not find any study that directly relates SHANK3 expression to AML clinical outcome or pathophysiology. This discovery may form the basis for future prospective clinical studies.

PRESTO comes as a compiled stand-alone application for Windows and Mac with an interface to direct users through the pre-processing, dimensionality reduction, and visualization steps. A full user guide is available on the project home page. PRESTO exports the gene groups, with their expression values, as a .csv file to be used in further downstream analysis. Future updates will include streamlining the process to re-perform PRESTO on a large identified gene group, to be able to narrow a gene signature in one step. The PRESTO graphical interface, along with a user manual, can be found on Github (https://github.com/saramcardle/PRESTO.)

In conclusion, PRESTO is powerful, intuitive, and insensitive to cohort size. It is particularly well suited to analyze large data sets with ordinary computer systems. PRESTO has opened a new window into -omics data across many samples and conditions, which are increasingly used in precision medicine studies.

## Implementation

### Algorithm Overview

PRESTO finds co-expression networks in -omics data from experiments with many samples, conditions, or time points (Fig. 1). The method includes pre-processing by thresholding on variability and expression, dimensionality reduction by tSNE, and visualization options to aid in interpretation. The algorithm is blinded to clinical information, enabling an unsupervised, hypothesis-free analysis of disease phenotypes in thousands of patients or conditions. Gene groups are exported for downstream analysis, functional annotation, diagnostic signatures, and survival predictions. These characteristics make PRESTO uniquely useful for the analysis of large clinical -omics datasets. The package comes as a stand-alone Matlab application (no Matlab license required) with an interactive user interface (Fig. 2a).

The input to PRESTO is any data matrix of microarray (log_2_-transformed and quantile normalized as robust multi-array average (RMA)), RNAseq (RPKM values, linear), or mass spectrometry data (spectral counts) from isolated cells or bulk biopsies. Markers (gene or protein expression) are input as rows and samples/conditions as columns. The minimum number of samples tested is 10, the maximum is 10,000 and 20-100 typically produce good results. The names of the markers and the samples/conditions are not used during the analysis, but are tracked, so that the same genes can be found in both conditions in the final result.

The first step in pre-processing is to remove genes with many values near or below the limit of detection to reduce the effect of noise and artificial minimums, as has been used previously in other applications^55, 56^. This threshold, and the number of samples which must meet that threshold, can be changed, even to 0 to remove this requirement entirely. It is important to remove any data points that have 0 (or the lower detection limit) expression in all samples. The data is further filtered for only those markers that show high variation between samples, based on the coefficient of variation (CoV) for each observation across samples^30^. Because variation is known to be a function of gene or protein expression^57^, ^58^, the data is split into deciles based on the average value for each observation across samples. The median CoV for each decile is calculated and the CoV threshold is set as a multiple of the median (Fig. 2b). This user-defined factor should be optimized for each data set. In our testing, a range of 1500 to 4000 genes is optimal for processing time and resolution, though the algorithm works with much larger or smaller data sets (range tested: 200 to 100,000). Performance analysis shows that the computing time changes negligibly with the number of columns (i.e. samples) and exponentially with the number of markers (i.e. genes, Fig. 2c). The expression values for each gene are normalized by dividing by the gene’s mean across all samples. This is to ensure that networks are identified based on their pattern across samples, not on their average expression. The resulting matrix of values after filtering and normalization are unitless, where each row (representing 1 gene or protein) has a mean value of 1 and a range from zero to the maximum of the dynamic range of the experimental method. Unlike the original tSNE publication, PCA was not found to accurately reduce the dimensionality of gene expression data^14^, so it was not used during the preprocessing step.

PRESTO uses t-Stochastic Neighbor Embedding (tSNE) scripts written as part of the Matlab dimensionality reduction package by the van der Maaten group that have been modified for this application. tSNE is a non-linear machine learning tool for detecting similarities between data points in high-dimensional data sets^19^. It calculates the distances between genes in the raw data with many samples, and then attempts to map those relationships into lower-dimensional space through gradient-descent optimization. The result is a 2D scatter plot where points (representing genes or proteins) that are close to each other have similar co-expression patterns across the samples in the input data set. There are two user-defined inputs into tSNE-“perplexity” and the number of iterations for optimization. “Perplexity” is ‘a smooth measure of the effective number of neighbors’^19^, which is loosely related to the size a discovered network. Typical values for gene expression data are 30-100. The iteration number determines how long it will spend trying to optimize the location of the points in 2D to best represent the multi-dimensional relationships. The original tSNE Matlab code has been modified to use the algorithm’s built-in cost function (a measure of how well the 2D dot placement represents the hyperdimensional distances) to monitor progress of the iterations. Optimization ends when the calculated cost stops decreasing, or when it reaches a minimum value of 0.2. This allows users to set a high number of iterations to ensure complete convergence without wasting time after results have stopped improving. Furthermore, random initial seed points for the 2D can be generated and saved. For paired analysis, it is highly recommended to generate seeds before starting tSNE. The points will be generated such that each gene starts in the same location for all groups or timepoints. This makes it likely that if a gene is co-regulated similarly between two conditions, the two dots representing that gene in both conditions will appear near each other in the final result.

The result is a set of X and Y coordinates (tSNE parameters 1 and 2) for each gene or protein that can be visualized as a scatter plot. The points appear to localize to multiple dense regions, representing genes that are expressed in similar patterns across the samples. Importantly, these spatial relationships are non-linear and non-deterministic, so distance between points on a scatter plot cannot be directly translated into multidimensional distance. Groups can be outlined manually based on apparent boundaries. Automated data clustering is an ongoing research area, but no universal “superior” algorithm has emerged yet^28^. The PRESTO user interface includes a module for DBScan^29^ for automatic clustering. The minimum number of points and epsilon value are user-defined. PRESTO exports the X and Y coordinates, as well as the expression values for each filtered gene to a spreadsheet.

A variety of visualization options are available as part of the PRESTO user interface to aid in biological interpretation of the results. It can show the marker expression of an individual sample or the average of all samples. PRESTO can also calculate a relative ranking of every sample for every marker. The sample that expresses that marker the least is assigned a value of 1, and the value increases successively for samples with greater expression (up to the number of samples). Then, the values for each marker for a particular sample can be displayed as a color coded scatter plot. The 2-dimensional data can be converted into a density map to more clearly delineate the boundaries between gates. If the samples are annotated in the input data with different group names, the expression values can be averaged for each group and the expression and rankings plot can be displayed by annotation name instead of sample name.

### Analysis of Matched Samples

Complex experimental designs involving biologically varying samples matched between multiple conditions (for instance, many individual patients sampled during multiple timepoints, or a range of mouse breeds stimulated with multiple treatments) can be used to analyze not just co-expression networks, but how those networks can be altered. PRESTO is able to analyze the co-expression networks within each condition or timepoint, and how those networks vary. The expression data for each condition/timepoint is entered as a separate matrix. For a gene to pass the pre-processing filters, the minimum expression threshold must be met in every condition (i.e., if a gene is not detected in one condition, it will not be analyzed at all). However, a gene needs only to pass the CoV filter in 1 of the conditions. Each gene will appear repeatedly in different rows in the final analyzed matrix (gene_name_condition1 and gene_name_condition2, etc), but in the same sample column.

After dimensionality reduction, the resulting scatter plot can be split to show the location of each gene in the different conditions. The location of the points can be directly compared to approximate how much the pattern of expression changes between the conditions. Groups can be defined on the graph of the overlay of all conditions or based only on one condition and then applied to the others. If a gene is located in nearly the same place in both graphs, it suggests that the pattern of expression across the samples is similar between the conditions. However, the reverse is true if the point moves to a different gate. This aids in understanding changes in co-expression networks between conditions.

## Methods

### Sensitivity Analysis of PRESTO

The sensitivity, robustness, and reproducibility of the PRESTO algorithm were tested using the human PBMC transcriptomes as well as the 75 strain HMDP LPS-treated data set. For the LPS-treated macrophages, the standard settings to which all changes were compared were: a minimum expression threshold of 1 sample with an RMA of at least 3, a variance cutoff of 1.5-fold median CoV per decile (2,423 genes selected), a “perplexity” of 50, and a particular random seeding. For the PBMCs, the standard settings to which all changes were compared were: a minimum expression threshold of 1 sample with an RMA of at least 0, a variance cutoff of 2-fold median CoV per decile (1,298 genes selected), a “perplexity” of 50, and a particular random seeding.

We generated graphs with different random seeds, “perplexity” of 10-750, or variance cutoffs of 0-2.5 fold median CoV (Supplemental Figs. 4 and 6). Additionally, we tested the effect of removal of 10% or 50% of the genes, or 1 or half of the samples (Supplemental Figs. 4 and 6). For the LPS data set, a vertical color gradient was applied to one graph (Repeat 1 on Supplemental Fig. 1), and then mapped onto other graphs to visualize differences in gene localization. For the PBMC data set the gate ID colors from the original data were mapped onto all other plots. The change in each case from the results of the standard settings were calculated with Jansen-Shannon Divergence, using the “compare_maps” Matlab script written by Pe’er, at al., as part of the Cyt package^27^. As a control, the Jansen-Shannon Divergence was calculated between the standard map and one of the same number of points that were randomly generated points within the same X and Y ranges.

### Microarray data

All analyses were performed on publicly available data sets. The details about the data sets, sample or patient characteristics, references, and PRESTO settings are listed in Supplemental Table 1. For all microarray data, probe sets were collapsed by taking the probe with the maximum expression for each gene, using the GenePattern 2.0 framework^59^.

For GSE12417 (AML test set), the maximum expression value between the two chips in Cohort 1 (GPL96 and GPL97) was used. Also, following Ng. et al^43^, we removed 3 patients whose sample came from peripheral blood instead of bone marrow or who were diagnosed with a disease other than AML.

### Comparative Bioinformatics Analysis

Hierarchical clustered heat maps were generated using the Broad Institute’s tool Morpheus. Average linking was used and the dendrogram was cut to best approximate the number of PRESTO gates. Principal component analysis was performed in Matlab. For some examples, a network of strongly co-expressed genes was constructed for the highly variable genes, using WGCNA^18^. Briefly, unsupervised transcriptional network for the highly variable genes was constructed by generating a signed co-expression network through creation of a matrix of Pearson correlations. This correlation matrix was used to calculate the adjacency matrix through soft thresholding by raising it to a power β (12). Based on the adjacency matrix, interconnectedness (topological overlap matrix) of each gene pair was computed^60^. Further, average linkage hierarchical clustering was done on the topological overlap matrix, and using the dynamic tree cut algorithm^61^ the branches were cut into defined modules. The obtained modules were compared with PRESTO patterns. DE expression analysis was performed on Gene Pattern using the Comparative Marker Selection toolkit using a two-sided t-test and a one-versus-all phenotype comparison. The significance threshold included FDR (Benjamini Hochberg) < 0.01 and p < 0.01. Genes with a fold change of 1.5 or greater were considered.

### Gene enrichment analysis

Gene set enrichment analysis (GSEA) determines whether a gene signature is significantly enriched in one of two conditions analyzed^62^. We performed GSEA on kidney biopsy transcriptomes of patients with antibody-mediated rejection compared to healthy controls (GSE36059, 20,647 genes) ^40^. The input gene signature was the list of all gate 1 genes in Figure 6 (GSE50084). The GSEA algorithm determines the mean gene expressions across all datasets/patients in both conditions, calculates the differential expression, and determines whether the individual genes of the signature are specific for one condition. GSEA standard settings were used (weighted, 100 iterations).

Bulk RNA deep deconvolution was performed using CIBERSORT^63^. We used the mean expression values of gate 1 genes across all patients with healthy transplants or rejected transplants as signature. As output CIBERSORT determines the estimated fraction of RNA in the sample explained by the signature (1 = full overlap, 0 = no detection). All enrichment p-values were < 0.001. Importantly, there is currently no p-value for detection limits, and a systematic error of over- or underrepresentation has been noted. For this reason, we also included biopsy transcriptomes of healthy nephrectomies, where no signature enrichment could be detected.

### Survival analysis

For some cases (Figure 6, Supplemental Figure 8), the impact of gene signatures on survival in published patient cohorts was determined using ProgGeneV2^64^. This tool facilitates patient survival stratification based on gene expression values. The cohort was divided by the median of the mean gene expression of the relevant network (all genes in one gate averaged as combined signature). The log-rank test and hazard ratio are indicated.

In other cases (Figure 7), gene signatures were condensed into a single score by defining weighting coefficients for each gene using Cox Proportional Hazard Regression^65^ (CPHR) in Matlab. The total hazard score for each patient was calculated, and patients were stratified as above or below the median hazard score for each signature. Kaplan-Meier^66^ curves were calculated in Prism, and the p-value and hazard ratio are shown.

### Functional Annotation

Each gene list was uploaded to DAVID^32^, ^67^ (version 6.8) for functional annotation. Annotation lists from each gene network were sorted by log(p-value) and relevant significant annotations were manually selected from the top 50 GO terms. Some gene lists were uploaded to Qiagen’s Ingenuity Pathway Analysis^68^ software. For proteomics data, the gene name for each identified protein was used. For each gene list, a ‘core analysis’ was performed, followed by a comparison analysis between major gates. Enrichment lists from the ‘Disease and Function’ feature were exported to Morpheus for heatmap visualization and hierarchical clustering – available through the Broad Institute (https://software.broadinstitute.org/GENE-E/index.html). Upstream regulators were determined using IPA with a p-value of overlap < 0.05.

### Statistics

Genes from each identified gate were averaged per sample, and the normality of the distribution was evaluated using D’Agostino-Pearson omnibus test. Samples were compared using either a non-paired two-tailed t-test or Mann-Whitney U test. In paired samples, a Wilcoxon matched-pairs signed rank test was performed. ^∗^ < 0.05, ^∗∗^ < 0.01, ^∗∗∗^ < 0.001. The three conditions of the RNA deconvolution analysis were compared using Kruskal-Wallis with Dunn’s correction for multiple comparisons.

## Footnotes

**Author contributions** SM developed the PRESTO algorithm and the Matlab interface. KB analyzed all clinical data. EE investigated cluster biology. ABP performed comparative analysis to published methods. KL conceived and supervised the study. SM, KB, EE, and KL wrote the manuscript.

## Acknowledgements

This work was funded by a La Jolla Institute for Allergy and Immunology institutional grant. K.B was supported by a Deutsche Forschungsgemeinschaft (DFG) grant (BU 3247_1).

